# HiTE: An accurate dynamic boundary adjustment approach for full-length Transposable Elements detection and annotation in Genome Assemblies

**DOI:** 10.1101/2023.05.23.541879

**Authors:** Kang Hu, Minghua Xu, You Zou, Jianxin Wang

## Abstract

Recent advancements in genome assembly have greatly improved the prospects for comprehensive annotation of Transposable Elements (TEs). However, existing methods for TE annotation using genome assemblies are less accurate and robust, requiring extensive manual editing. In addition, the currently available gold-standard TE databases are not comprehensive, even for extensively studied species, highlighting the critical need for an automated TE detection method to supplement existing repositories. In this study, we introduce HiTE, an accurate dynamic boundary adjustment approach designed to detect full-length TEs. The experimental results demonstrate that HiTE identified 932 perfect TE models with a precision of 0.971 on the rice reference genome, which are 142% and 4.42% higher than the state-of-the-art tool RepeatModeler2, respectively. Furthermore, HiTE discovers over 800 novel TIR elements with well-defined structures that are not included in known libraries, enabling the discovery of new insights. We have also implemented a Nextflow version of HiTE to enhance its parallelism reproducibility, and portability.

## Introduction

Transposable elements (TEs), which make up the majority of repetitive regions in most eukaryotic species^1–3^, are known to have a significant impact on genome evolution and intraspecific genomic diversity^4, 5^. TEs have been found to play key role in human diseases and crop breeding by interrupting or regulating the key genes^6–8^. Identifying intact TEs is challenging due to various complications^9^, including but not limited to: (i) the varying degradation rates of TEs, which can lead to the loss of structural signals^10^; (ii) the complex pattern of TE sequences, resulting from random deletions, insertions^11, 12^, and nested TE^13^; (iii) the difficulty to determining the true ends of highly fragmented TE instances^14^; (iv) the obstacle to constructing full-length TE models posed by the abundance of fragmented TEs; (v) the confounding impact of regional homology between unrelated TEs on their identification and classification; and (vi) the risk of erroneously identifying high copy numbers of segmental duplications or tandem repeats as putative TE instances.

There are several tools available for the automated identification and annotation of TEs, which can be broadly divided into three categories: (i) De novo methods, (ii) Signature-based methods, and (iii) TE discovery pipelines. By identifying exact or closely matching repetitions, de novo methods can identify novel TE instances that do not belong to a known family of TE, which mainly includes a (spaced) k-mer-based or self-comparison approach. While k-mer-based approaches, such as RepeatScout^15^ and P-Clouds^16^, are better suited for dealing with young TEs with plenty of copies, they may produce highly fragmented sequences for older TEs with diverse or complex patterns. On the other hand, self-comparison methods, such as Grouper^17^, RECON^18^, and PILER^19^, can identify more sophisticated TE families using intensive and sensitive alignments, but accurately clustering highly fragmented and mosaic TE sequences remains challenging. Signature-based methods, such as LTRharvest^20^, LTR_retriever^21^, Generic Repeat Finder^22^, EAHelitron^23^, HelitronScanner^24^, and MITE-Hunter^25^, identify TEs based on family-specific features. These methods can overcome the limitations of purely de novo methods that may miss well-characterized TEs with low copy numbers. However, signature-based methods are prone to false positives due to the weak structural characteristics of many TEs. TE discovery pipelines, like EDTA^26^ and RepeatModeler2^27^, combine different TE identification tools to comprehensively identify all types of TEs within a given genome. While these pipelines can overcome the limitations of individual tools, they also introduce their inherent defects and require careful handling of redundant results. A high-quality TE library can help overcome these challenges by providing a comprehensive and structured collection of TEs. After years of manual curation, Repbase^28^ and Dfam^29^ are high-quality consensus libraries for a limited set of species, while all automatically generated TE libraries still require extensive manual editing^30^.

An ideal TE library should only contain full-length TE models^9^. However, the reality is that almost all tools will inevitably introduce some fragmented TE models. RepeatModeler2 introduces a benchmarking method to quantify full-length and fragmented TE models. Specifically, it uses *Perfect* indicators to represent full-length TE models, while *Good* and *Present* indicators represent fragmented TE models. In this study, we introduce HiTE, an accurate dynamic boundary adjustment method for detecting full-length TEs with high precision. Using highly conservative structural features and multiple copies, HiTE discovers many known and novel TE instances and produces a high-quality, structurally intact, and classified TE library. We further develop a Nextflow^31^ pipeline of HiTE, which enhances the reproducibility and portability of HiTE. Comprehensive experiments on four model species, ranging from plants to animals, show that HiTE outperforms other tools in terms of accuracy and the identification of full-length TE models. The results of this study demonstrate the potential of HiTE to the creation of a high-quality TE library for a wide range of species.

## Results

### Overview of HiTE

Purely de novo methods for detecting TEs based on sequence repetition alone may miss low-copy but well-characterized instances, whereas signature-based methods are susceptible to false positives, owing to the poor structural characteristics of certain TEs^9, 32^. HiTE is an automated TE annotation pipeline that combines the strengths of de novo and signature-based methods, aiming to produce a high-quality TE library. To achieve comprehensive TE annotation, HiTE employs various useful tools and does not require manual intervention. Considering the diverse structural characteristics and distribution of TEs in the genome, HiTE develops three modules, namely de novo TE searching, structural-based LTR (long terminal repeat) searching, and homology-based non-LTR searching (Fig. 1c, h, and i). These modules can identify nearly all types of transposons, including LTR, TIR (terminal inverted repeat), Helitron, LINE (long interspersed nuclear element), and SINE (short interspersed nuclear element)^33^.

Firstly, HiTE divides the genome assembly into smaller chunks to reduce computation during a single run (Fig. 1b). Secondly, we have devised HiTE-FMEA, a module that uses sensitive alignment algorithms to identify TE models with coarse-grained boundaries (Fig. 1c, Methods). Then, we have developed HiTE-TIR and HiTE-Helitron modules to identify TE models with fine-grained boundaries (Fig. 1e, Methods). To minimize false positives, such as tandem repeats and segmental duplications, we have also implemented multiple filtering methods, including the removal of candidates containing extensive tandem sequences or significant homology outside their copy boundaries (Fig. 1d and f, Methods). Next, we use LTR_FINDER^34^ and LTRharvest^20^ to identify all candidate LTR-RTs (LTR retrotransposons), which are then subjected to stringent filtering using LTR_retriever^21^ to identify reliable LTR-RTs (Fig. 1h). In addition, we have developed a homology-based non-LTR searching module to achieve high-precision non-LTR annotation (Fig. 1i). Finally, we collect all types of TEs together to generate an intact TE consensus library and implement a parallel version of RepeatClassifier, a component of RepeatModeler2^27^, to speed up the classification process without compromising accuracy (Fig. 1g, Methods).

**Fig. 1.**
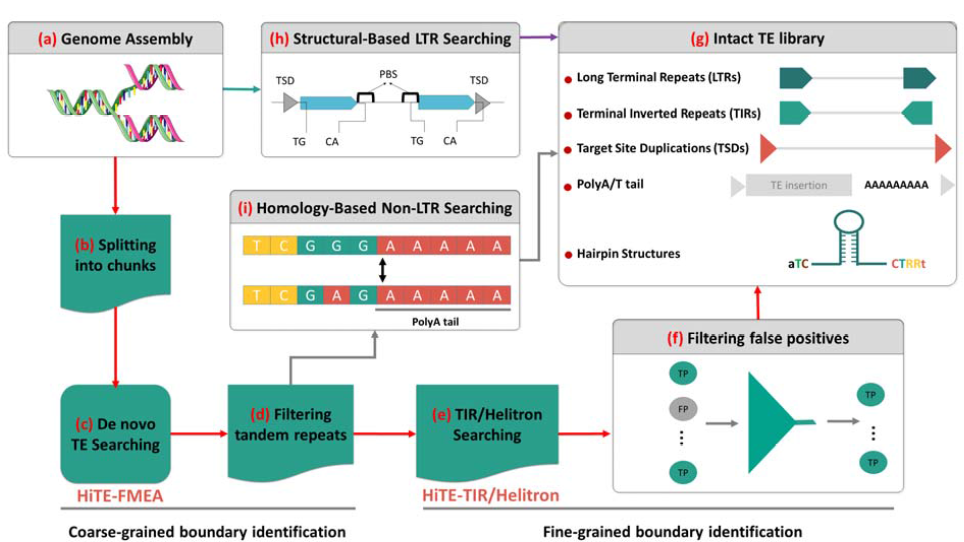
Workflow of the HiTE pipeline for TE annotation **a** HiTE takes the genome assembly as input. **b** The genome assembly is split into chunks to reduce the amount of single-round computation. **c, h, and i** Three main modules of HiTE are developed to identify different types of TEs. **d** Sequences containing a large proportion of tandem repeats are filtered out as false positives. **e** Signature-based methods are used to identify TIR and Helitron elements with fine-grained boundaries. **f** False positives are filtered out using reliable strategies, such as filtering candidates with homology beyond their copy boundaries. g TE libraries generated by HiTE have intact TE structures.

### HiTE accurately detects more intact TE models

To evaluate the performance of different TE identification methods, we use the benchmarking methods released in two recent studies, RepeatModeler2 and EDTA^26^ (BM_RM2 and BM_EDTA hereafter). BM_RM2 is based on RepeatMasker (4.1.1) and a custom bash script (https://github.com/jmf422/TE_annotation/blob/master/get_family_summary_paper.sh) provided by RepeatModeler2. BM_EDTA is based on the Perl script "lib-test.pl" included in EDTA. TE models are considered *Perfect* matches with >95% sequence similarity and >95% length coverage for a family consensus in the gold standard library. An ideal TE library should only contain full-length TE models, which can be evaluated by the *Perfect* indicator from BM_RM2. Given that the protein sequence required for transposition is contained within the full-length TE, the quantity of *Perfect* models is the most meaningful metric for evaluating TE integrity and biological significance^27, 35^. It is worth noting that a full-length TE sequence spanning 1 kbp holds more biological significance than ten individual fragments of 100 bp, since the former may contain the complete TE protein structure, which is indispensable for TE transposition. Since errors in the library can propagate throughout the whole-genome annotation process^36^, it is also essential to account for the rate of false positives in the TE library. To address this challenge, we use the *Precision* indicator from BM_EDTA to estimate false positives. By using both the *Perfect* indicator from BM_RM2 and the *Precision* indicator from BM_EDTA, we can evaluate the TE library generated by various tools in terms of precision and the integrity of TE models.

As shown in Fig. 2a, HiTE outperforms the other three tools on all datasets. HiTE restores 932, 69, 625, and 95 full-length (*Perfect*) TE models on O. sativa, D. melanogaster, D. rerio, and C. briggsae, which are significantly higher than the other three tools, respectively. At the same time, more *Good* and *Present* TE models always mean more fragmented sequences, which can hinder the identification and classification of full-length TE models. HiTE also discovers relatively fewer fragmented TEs than other tools. We noticed that RepeatScout and RepeatModeler2 both achieved consistently high sensitivity using BM_EDTA, which is also validated by EDTA^26^. However, the evaluation of BM_EDTA relies on the count of matched bases instead of the complete TE sequence, potentially leading to the inclusion of short false-positive sequences or fragments, which in turn can cause an erroneous elevation of sensitivity. As demonstrated in the cases of O. sativa and D. melanogaster, RepeatScout generated TE models with lower structural integrity but maintained high sensitivity levels (Fig. 2). Similarly, the HiTE-FMEA module of HiTE exhibited high sensitivity but also included a significant number of fragmented and false-positive sequences (Fig. 3b).

To reduce false-positive sequences and fragmented TEs, HiTE uses multiple filtering methods to ensure that the candidates discovered have complete TE structures (Methods). In summary, HiTE gets 0.8082-0.9776 precisions, which are higher than the other three tools on all four datasets (Fig. 2b). The sensitivity of HiTE is relatively low due to the exclusion of numerous fragmented and false positive sequences that lack clear structures or multiple-copy support. The inclusion of such sequences would increase the fragmentation of the TE library, thereby impeding subsequent analyses, such as TE annotation. HiTE outperforms all evaluated methods in terms of precision and produces the highest number of *Perfect* TE models. The results demonstrate that HiTE can accurately detect full-length TEs models.

**Fig. 2.**
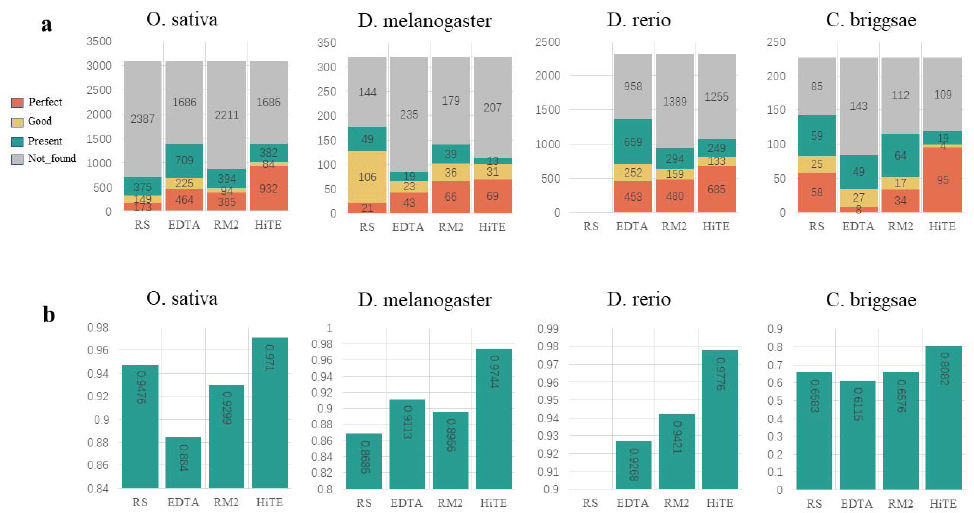
Evaluation of HiTE on structurally intact and high-precision TE identification using four different species. a The number of TE models identified using BM_RM2. b The precision comparison based on BM_EDTA. Since RepeatScout cannot process genomes larger than 1 GB, it has no results for D. rerio. RS: RepeatScout; RM2: RepeatModeler2.

### HiTE-FMEA detects TE models with coarse-grained boundaries

De novo methods based on identifying exact or closely matching repetitions can be used to discover novel instances that do not belong to known TE families. However, the generation of fragmented TEs is a common problem in these methods, resulting in many misclassifications. To address this issue, a fault-tolerant mapping expansion algorithm of HiTE (HiTE-FMEA) was developed to cross large gaps caused by insertion, deletion, and nested TEs, while preserving the integrity of the TE structures (Fig. 3a, Methods). Nevertheless, HiTE-FMEA can only identify TE models with coarse-grained boundaries, leading to fragments and ambiguous ends of candidates. For instance, HiTE-FMEA achieved high sensitivity of 0.9753 but showed suboptimal performance in terms of specificity (0.8615), accuracy (0.9171), and precision (0.8707), which suggests the presence of false-positive fragments (Fig. 3b).

**Fig. 3.**
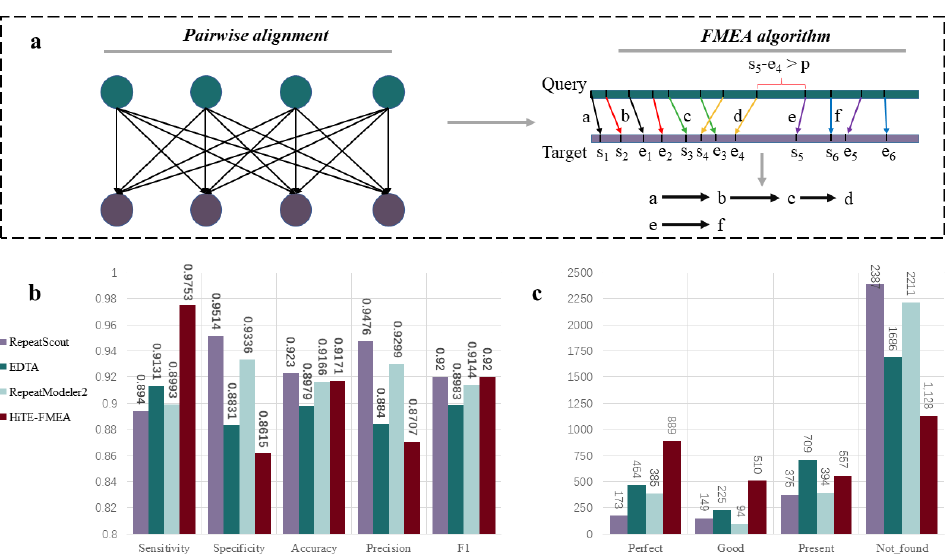
Evaluation of HiTE-FMEA for TE annotation based on O. sativa. **a** Illustration of pairwise alignments of genome assembly contigs and the fault-tolerant mapping expansion algorithm, FMEA. Two TE instances included in a contig sequence, a-d and e-f, are identified by FMEA. **b, c** Evaluation of HiTE-FMEA against RepeatScout, EDTA, and RepeatModeler2 using BM_EDTA and BM_RM2, respectively.

Upon manual inspection, the sequences identified by HiTE-FMEA have been found to include not only TE elements but also non-TE elements, such as tandem repeats, segmental duplications, and multi-copy genes. While HiTE-FMEA may not be able to discover the exact true ends of TE instances, it has been demonstrated to restore the most *Perfect* TE models when compared to the other three methods. Specifically, HiTE-FMEA has identified 889 *Perfect* TE models, which is considerably higher (by 414%, 92%, and 131%) than the corresponding models identified by RepeatScout, EDTA, and RepeatModeler2, respectively (Fig. 3c). However, HiTE-FMEA has also identified 510 *Good* and 557 *Present* TE models, suggesting the existence of fragmented TE sequences that require further identification of precise boundaries.

### HiTE-TIR/Helitron detects TE models with fine-grained boundaries

HiTE-FMEA shows limitations in accurately identifying TEs. At the same time, identifying class II transposable elements^33^, such as Terminal inverted repeat (TIR)^37^ and Helitron elements^38^, is challenging due to their weak structural signals. To address this issue, we design structural-based identification methods for TIR and Helitron elements, called HiTE-TIR/Helitron (Fig. 4a, Methods), which can produce high-precision and structurally intact TE models. To avoid the propagation of errors in the annotation library throughout the whole-genome annotation process, HiTE prioritizes higher precision at the cost of some sensitivity. We design a false-positive filtering method (Fig. 4a, Methods) to reduce false positives and generate a clean library. As shown in Fig. 4b, by using these methods, HiTE achieves 0.971 precision, which are significantly higher than HiTE-FMEA. Due to the presence of numerous fragmented and false positive sequences, HiTE-FMEA generated a large library of 147 Mb. In contrast, HiTE accurately identified TE boundaries and filtered out a significant number of fragmented sequences and false positives, resulting in a more concise library of 7.3 Mb. For example, HiTE significantly increases the number of identified *Perfect* TE models compared to HiTE-FMEA, while reducing the number of fragmented sequences represented by *Good* and *Present* TE models (Fig. 4c). HiTE identifies longer TE models than HiTE-FMEA, with a median length of 994 bp compared to 399 bp for HiTE-FMEA. These results demonstrate that the fine-grained TE boundary detection can improve the accuracy of TE identification and reduce false positives.

**Fig. 4.**
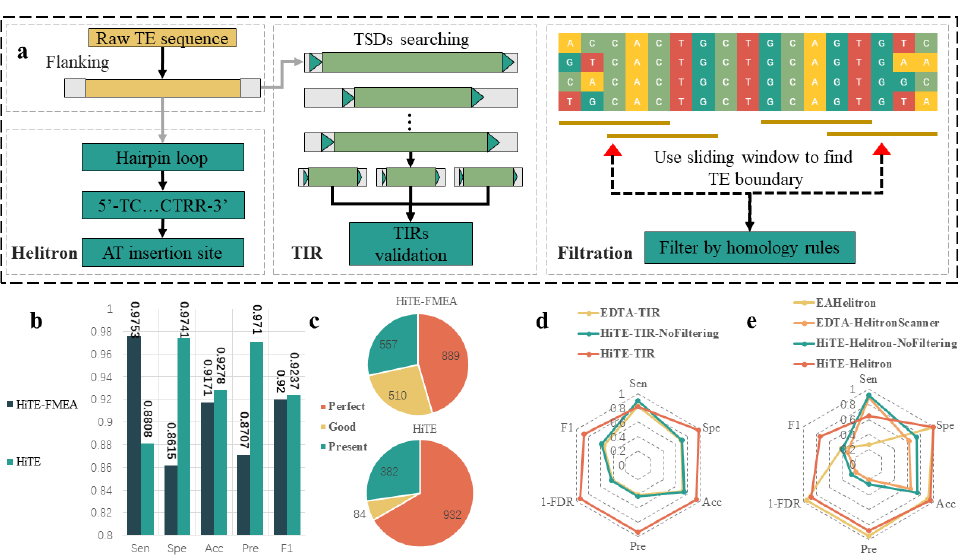
Identification of real TEs with fine-grained TE boundaries based on O. sativa. **a** Pipeline of HiTE for identifying real TIR and Helitron elements. Searching for confident structures reveals the true ends of TIR and Helitron elements. The robust evidence of true TEs is checked by non-homology flanking regions of multiple copies. **b, c** Evaluation of HiTE against HiTE-FMEA using BM_EDTA and BM_RM2, respectively. **d** Evaluation of HiTE-TIR against HiTE-TIR-NoFiltering and EDTA-TIR using BM_EDTA. **e** Evaluation of HiTE-Helitron against HiTE-Helitron-NoFiltering, EDTA-HelitronScanner, and EAHelitron using BM_EDTA. Sen: Sensitivity; Spe: Specificity; Acc: Accuracy; Pre: Precision; FDR: False discovery rate.

We evaluate the performance of the false-positive filtration module on the identification of TIR and Helitron elements. As shown in Figs. 4d and e, HiTE without false-positive filtration (HiTE-xxx-NoFiltering) achieves similar performance with EDTA (EDTA-xxx). HiTE-TIR gets a precision of 0.9312 and an F1 score of 0.87, which are 124% and 57% higher than EDTA-TIR (Fig. 4d). At the same time, HiTE-TIR identifies longer TE models and produces smaller library than HiTE-TIR-NoFiltering.

HelitronScanner and EAHelitron are the only available tools for identifying reliable Helitrons. Although EDTA can increase accuracy and specificity by refining HelitronScanner results^26^, its ability to precisely identify Helitrons is still limited, as shown in Fig. 4e. The evaluation of EAHelitron revealed that it had the highest precision but the lowest sensitivity and could not recover any gold standard models (Fig. 4e). To address this issue, we developed HiTE-Helitron by combining the FMEA algorithm and HelitronScanner. HiTE-Helitron achieves the best overall performance in identifying Helitrons with a precision of 0.8806 and an F1 score of 0.7432, which are 344% and 129% higher than EDTA-HelitronScanner, respectively (Fig. 4e). Moreover, HiTE-Helitron identifies longer TE models and generates a considerably smaller library in comparison to HiTE-Helitron-NoFiltering. The experimental results demonstrate that the fine-grained TE boundary detection significantly improves overall performance.

### HiTE discovers known characteristics of TEs

As an additional assessment of the ability for HiTE to discover known TEs, we run RepeatMasker^39^ with each output library generated by different tools and measure the percentage of the genome masked by each major TE subclass (Fig. 5a). HiTE restores the TE landscapes of these species consistent with the curated libraries. The genome of O. sativa is known to contain DNA-TIR and LTR elements in close proportions^40^, which is recovered by our HiTE library. Our results show that the genome of D. melanogaster is dominated by retrotransposons, especially LTR and LINE retroelements^41^. The genome of D. rerio is dominated by class II DNA-TIR transposons, but it also has a diverse composition of LTR retroelements with many distinct families^42^. The HiTE library discovered an abundant percentage of DNA-TIR elements in the genome of C. briggsae, which achieves a similar proportion to the curated library^43^.

**Fig. 5.**
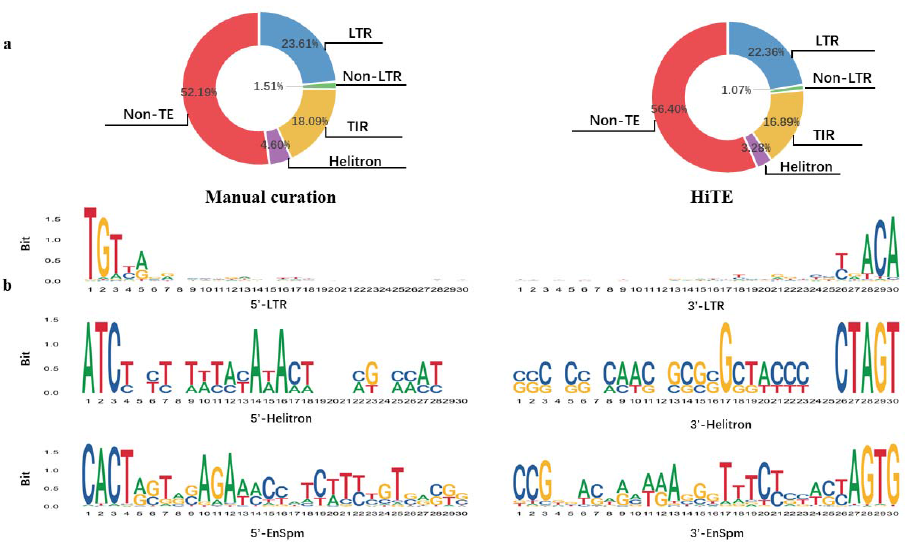
Evaluation of HiTE on discovering the landscape and structural features of TEs. **a** Percentage of the rice genome masked by each major subclass using TE libraries generated from manual curation and HiTE. **b** Sequence logos of terminal sequences of LTR, Helitron, and EnSpm candidates discovered by HiTE. Candidates at the 5’ ends with the starting position labeled as 1, and candidates at the 3’ end with the last position labeled as 30.

To validate the precision of the transposons identified by HiTE, we performed an analysis of the terminal sequences of LTR, Helitron, and EnSpm elements by generating sequence logos that indicate the nucleotide usage at each position in their terminals (Fig. 5b). Our findings are consistent with prior research, which reveals that LTR elements are generally flanked by 2-bp palindromic motifs, commonly 5’-TG…CA-3’^44^. We also evaluated the insertion times of two types of LTR-RTs, Copia and Gypsy, in rice and maize, and the results were consistent with previous literature^45, 46^. We observed that Helitron elements are inserted into an AT target site and have no terminal inverted repeats, with the canonical terminal structure of 5’-TC…CTRR-3’ (where 5’-TC…CTAG-3’ predominates). We also noted a higher AT content at the 5’ ends and an enriched CG content at the 3’ terminal, particularly at the 1 and 17 positions (Fig. 5b), which contribute to a canonical Helitron feature, such as a hairpin loop, as previously reported^47, 48^. Furthermore, we identified highly conserved CACT(A/G) motifs in EnSpm short terminals, consistent with previous reports^49^. These results further confirm the accuracy and reliability of HiTE in identifying TEs.

## Discussion

The rapid advancement of sequencing technologies has led to more reliable genome assemblies, which gives a bright future to comprehensive annotation of TEs. However, inaccurate TE identification tools can produce TE libraries containing many errors, which can propagate throughout the whole-genome annotation process. In this study, we developed and validated HiTE, an accurate dynamic boundary adjustment method for detecting intact TEs. We developed multiple algorithms that make full use of the repetitive nature, conserved motifs, and structural features of TEs for accurate detection. We have demonstrated that HiTE can identify a significant number of novel and genuine transposons, which can serve as a valuable complement to the currently gold standard database.

Given that the protein sequence required for transposition is contained within the full-length TE, the quantity of full-length TE models is the most meaningful metric for evaluating TE integrity and biological significance. A key feature of HiTE is its capability to identify full-length TEs, even in the presence of large gaps caused by insertion, deletion, and nested TEs. Accurately determining the boundaries of TEs poses a significant challenge for automated methods, frequently requiring extensive manual identification and correction. Through gradually discovering the full-length TEs from coarse to fine-grained boundaries, HiTE can identify the precise boundaries of TEs and significantly reduce the need for manual intervention. By benchmarking on four model species with different TE landscapes, we demonstrated that HiTE outperforms other TE identification methods, including RepeatModeler2, EDTA, and RepeatScout, in identifying intact TE insertions. Additionally, HiTE achieves superior performance in identifying specific type of TEs, such as TIR and Helitron elements.

Currently, methods that use genome assembly for TE identification, including HiTE, face certain limitations. One of the major challenges is the weak structural characteristics of certain types of TEs, which can lead to the inclusion of false-positive sequences. Although we have greatly improved the identification performance of TIR and Helitron elements, there is still potential for improvement. For example, incorporating more comprehensive hairpin loop patterns in Helitron identification tools could significantly improve their performance. Another limitation is the potential for loss of genuine TEs in Repbase. To reduce false positives, TE candidates with divergent terminals or TSDs, or accidental homology outside the boundaries, are considered false positives and discarded, which may result in the loss of some real TE instances.

In summary, through the utilization of sensitive alignment algorithms and structural feature recognition, HiTE has demonstrated considerable potential as a reliable tool for TE identification and annotation using genome assembly. We anticipate that the methods proposed in this study will contribute to the analysis of genome variation research, with potential applications in areas such as human disease and crop breeding.

## Methods

### Data preparation

During the evaluation of HiTE, four model species, namely Oryza sativa (assembly IRGSP-1.0), Caenorhabditis briggsae (assembly CB4), Drosophila melanogaster (assembly Release 6 plus ISO1 MT), and Danio rerio (assembly GRCz11), were used as reference genomes. The curated libraries were obtained from RepBase26.05, and the appropriate parameters were used to generate TE libraries of RepeatScout, RepeatModeler2, and EDTA. A non-LTR library was generated by extracting known LINEs and SINEs from the Dfam library of RepeatMasker version 4.1.1 (http://www.repeatmasker.org), which is a public TE database available under the Creative Commons Zero (CC0) license. Additionally, another rice genome assembly (Oryza sativa L. ssp. japonica cv. “Nipponbare” v. MSU7)^50^ was utilized to verify that HiTE can detect a stowaway-like MITE (sMITE) closely related to the key gene Ghd2.

### Fault-tolerant mapping expansion algorithm of HiTE (HiTE-FMEA)

The identification of a single TE instance as multiple fragments can significantly hinder the accurate identification and classification of complete TE families^51, 52^. To overcome this issue, we developed a fault-tolerant mapping expansion algorithm, termed FMEA, which can span large gaps and identify the coarse-grained boundaries of TEs. The process of FMEA involves the following steps (Fig. 3a):

1. Sensitive pairwise alignment. We use BLASTN^53^ for pairwise alignment of the genome assembly sequences. To speed up this process, assembly sequences are segmented into smaller fragments of 1 Mb using the "--chrom_seg_length" option, and concurrent Python package is utilized for parallel acceleration^54^.
2. Fault-tolerant expansion. For each query sequence, adjacent alignments are first gathered based on their alignment positions on the target sequences, which are then sorted ascendingly for clustering. Next, within each cluster, the alignment is expanded if the next alignment is in its adjacent area, spanning alignment gaps caused by TE variations. As shown in Fig. 3a, two TE instances included in the query sequence, *a-d* and *e-f,* are identified by FMEA. Finally, each query sequence could align to multiple target sequences and generate multiple candidate instances, which are subsequently screened for redundancy. The longest sequences are taken, which usually contain one or more repeats.

### Structural-based TE searching of HiTE

We developed structural-based methods for TE identification, which exploit the characteristic features of TEs, such as long terminal repeats (LTRs) and terminal inverted repeats (TIRs) at both ends of LTR and TIR elements. Additionally, TEs are typically inserted into the genome with the formation of two short TSDs resulting from the repair of DNA double-strand breaks at the integration site^55^. The size of the TSDs can be used as a diagnostic feature for TE identification and classification. The structural-based TE searching methods of HiTE mainly include the following parts:

1. The identification of LTR retrotransposons (LTR-RTs). LTR-RTs are facilitated by their distinctive structural characteristics, including long direct repeat sequences ranging from 85 to 5000 bp, 2-bp palindromic motifs (5’-TG…CA-3’) at both ends, and 4-6 bp TSDs flanked at the insertion site (Fig. 1h). To identify LTR-RTs, we employed LTR_harvest and the parallel version of LTR_Finder^34, 56^ with "-seed 20 -minlenltr 100 -maxlenltr 7000 -similar 85 -motif TGCA -mintsd 4-maxtsd 6 -vic 10" and default parameters, respectively. We then use LTR_retriever as a stringent filtering method for the candidate LTR-RTs.
2. The identification of TIR elements. TIR elements have terminal inverted repeat sequences (usually a few bp to hundreds of bp) and conserved motif characteristics of specific superfamilies. Due to their short termini, TIR elements can be difficult to identify. To discover the structurally intact TIR elements, we first use the FMEA algorithms to find the coarse-grained boundaries of candidate TEs, which are then extended by a certain length to search for all legal TSDs (Fig. 4a). To reduce false positives, we identify TSDs that are identical, allowing for at most a 1-bp mismatch in those longer than 8 bp. Next, we use the itrsearch tool, included in TE Finder 2.30 (https://github.com/urgi-anagen/TE_finder), with the parameter "-i 0.7 - l 7" to search for TIRs. Since a sequence with a coarse-grained boundary may produce multiple candidates with legal TIRs and TSDs, we select the candidate that is closest to the coarse-grained boundaries as the TIR-like elements. Finally, the TIR-like elements are passed to the false-positive filtering module of HiTE to search for homology boundaries, adjust the true ends of TEs, and obtain confident TIR elements.
3. The identification of Helitron elements. Helitrons are a class of TEs that replicate through the rolling circle mechanism, and their weak structural signals make them particularly challenging to identify. To identify Helitrons, we first use HelitronScanner on the candidates with coarse-grained boundaries and search for the hairpin structures (Fig. 4a). Then, we select the candidates with complete Helitron structures flanked by 5’-A and 3’-T that are closest to the coarse-grained boundaries as the Helitron-like elements. Finally, the Helitron-like elements are passed to the false-positive filtering module of HiTE to search for homology boundaries, adjust the true ends of TEs, and obtain confident Helitrons.

### False-positive filtering of HiTE

False positives in the library can propagate throughout the whole-genome annotation processes. We have identified four major types of false positives that significantly affect the accuracy of TE identification:

1. Sequencing gaps. To reduce the impact of misassembles in repetitive sequences, we exclude any candidate TE sequences that contain contiguous gaps longer than 10 bp. These gaps represent the areas of highest uncertainty in a genome assembly and are more prone to misassembly errors.
2. Tandem repeat. We use Tandem Repeats Finder (TRF)^57^ to identify tandem repeats with parameters "2 7 7 80 10 50 500 -f -d -m". Sequences in which tandem repeats account for more than 50% of the whole sequence are filtered out. Furthermore, TIR candidates with tandem repeats exceeding 10 bp at the beginning of their terminals are also eliminated to avoid fake TIR elements, such as those that begin with ‘TA’ and end with ‘AT’, which are frequently associated with simple tandem repeats.
3. Spurious TIR elements with LTR terminals. The occurrence of short TIR terminals with legal TSDs within the LTR may generate spurious TIR candidates, leading to false positive results. To mitigate such outcomes, TIR candidates are aligned to the LTR-RTs identified by the LTR searching module of HiTE. TIR candidates with more than 80% overlap regions with LTR-RTs are regarded as false positives and subsequently removed from further analysis.
4. Candidates with homology beyond their copy boundaries. Many false positive sequences may possess TE-like structures by chance, leading to a significant amount of misclassification. To address this issue, we have developed a filtering method based on two principles, as depicted in Fig. 4a. Firstly, transposons should occur at least twice in the genome, disregarding old TEs that have evolved and diverged significantly over time. Secondly, the region outside the boundaries of TEs should be composed of random sequences with no homology. Therefore, we first extended both ends of the candidate TE copies and performed multiple sequence alignment between them. Subsequently, we used a 10-bp sliding window to detect homology boundaries in the multiple sequence alignment files. We then evaluated whether the candidate transposon identified by the homologous boundary met a specific transposon structure, such as the presence of a TSD and TIR structure for TIR transposons and a 5’-ATC…CTRR-3’ structure for Helitron transposons. Ultimately, we obtained a set of genuine TEs with well-defined structures.

### Homology-based non-LTR searching of HiTE

The identification of non-LTR elements, such as LINEs and SINEs, is challenging due to their variability and lack of discernible structural signals. To date, there are no structure-based methods that can accurately and comprehensively identify LINEs and SINEs^58, 59^. To address this issue, we have developed a high-precision annotation approach for non-LTR elements by creating a non-LTR library, which includes known LINEs and SINEs from the Dfam library of RepeatMasker version 4.1.1. This library is then used to search against the genome to identify confident non-LTR elements.

### Unwrapping nested TEs

Nested TEs, which are transposons inserted into other transposons, can complicate the identification of individual TE sequences. To address this issue, HiTE uses a three-step process to unwrap the nested TEs. Firstly, HiTE removes the full-length TEs contained in other sequences with more than 95% coverage and 95% identity and connects the remaining sequences. Secondly, sequences shorter than 100 bp are filtered out, while the remaining sequences are treated as new TE sequences. Finally, the process is iterated several times to unwrap heavily nested TEs. This process allows for the accurate identification and annotation of individual TE sequences even in the presence of nested TEs.

### Generation of a classified TE library

HiTE uses CD-HIT with the parameter "-aS 0.95 -aL 0.95 -c 0.8 -G 0 -g 1 -A 80" to generate consensus models of TEs. Before clustering, LTR-RTs are divided into 5’ LTRs, 3’ LTRs, and LTR internal regions. In addition, we implemented a parallel version of RepeatClassifier, a component of RepeatModeler2, to speed up the classification process without compromising accuracy.

Many TEs contain open reading frames (ORFs) that encode proteins necessary for their transposition. Predicting conserved protein domains in TEs can help with their classification into the different superfamilies. We performed the homology search between protein database and TE models using BLASTX and filtered out bad hits with e-value cut off 1e-20. The protein database of known TE peptides can be downloaded from the RepeatMasker. In addition, alignment results are often fragmented, making it difficult to discover the intact protein domains of TEs. To address this issue, we connected and bridged the fragmented alignments to create a more continuous domain area within the TE models. Finally, we generated a table that maps TE models to the corresponding protein domain locations, providing a convenient description of the protein structure of TEs.

### Evaluation method of HiTE

To evaluate the performance of HiTE and compare it with other TE detection tools, two benchmarking methods released in recent studies, RepeatModeler2 and EDTA, are employed.

1. Evaluation by benchmarking method of RepeatModeler2 (BM_RM2). BM_RM2 performs an alignment of the tested TE library with a gold standard library and categorizes the resulting matches into four levels: “Perfect”, “Good”, “Present”, and “Not found”.
2. Evaluation by benchmarking method of EDTA (BM_EDTA). BM_EDTA evaluates the performance of various tools by annotating the genome with the gold standard TE library and the tested TE library, which is generated by different tools. The assessment is conducted based on a summary of the total number of genomic DNA bases. Six metrics comprising sensitivity, specificity, accuracy, precision, FDR, and F1, are used to characterize the annotation performance of the tested library. BM_EDTA can display false-positive rates, and BM_RM2 aims to reflect the integrity of TE models. By using both the *Perfect* indicator from BM_RM2 and the *Precision* indicator from BM_EDTA, we can evaluate the TE library generated by various tools in terms of precision and the integrity of TE models.

## Data availability

The NCBI hosts reference genomes for five species [https://www.ncbi.nlm.nih.gov/], including Oryza sativa (assembly IRGSP-1.0), Caenorhabditis briggsae (assembly CB4), Drosophila melanogaster (assembly Release 6 plus ISO1 MT), Danio rerio (assembly GRCz11), and Zea mays (assembly Zm-B73-REFERENCE-NAM-5.0). The other rice genome (Oryza sativa L. ssp. Japonica cv. "Nipponbare" v. MSU7), as well as its annotation with respect to both genes and repeats, can be accessed through the Rice Genome Annotation Project [http://rice.uga.edu/]. Curated TE libraries can be found at Repbase [https://www.girinst.org/repbase/]. Additionally, TE libraries produced by HiTE, RepeatModeler2, EDTA, and RepeatScout, as well as novel TIR elements of rice discovered by HiTE, are publicly available in the GitHub repository CSU-KangHu/TE_annotation [https://github.com/CSU-KangHu/TE_annotation].

## Code availability

HiTE is publicly available at GitHub [https://github.com/CSU-KangHu/HiTE].

## Acknowledgments

This work was supported in part by the National Natural Science Foundation of China under Grants (Nos. 62150048), 111 Project (No. B18059), Fundamental Research Funds for the Central Universities of Central South University (2021zzts0208). This work was carried out in part using computing resources at the High Performance Computing Center of Central South University.

## Author contributions

J.X.W. conceived and designed this project. K.H. and J.X.W. conceived, designed, and implemented the HiTE. K.H. and X.M.H conducted the analyses. Y.Z. helped evaluate HiTE on the HPC cluster. K.H. and J.X.W. wrote the paper. All authors have read and approved the final version of this paper.

